# Genome-wide analysis of cell cycle-regulating genes in the symbiotic dinoflagellate *Symbiodinium minutum*

**DOI:** 10.1101/307546

**Authors:** Michael L. Cato, Hallie D. Jester, Adam Lavertu, Audrey Lyman, Lacey M. Tallent, Geoffrey C. Mitchell

## Abstract

A critical relationship exists between reef-building corals and their photosynthetic endosymbionts. As important as this relationship is for reef health, it is exquisitely delicate—exposure to temperatures only marginally above the average summer maximum can cause corals to bleach, expelling their resident algae. Interestingly, several studies indicate that failure of corals to properly regulate symbiont cell divisions at high temperatures may cause bleaching. This needs to be further investigated, but first, it is necessary to decipher the molecular mechanisms controlingl the cell division cycle in these organisms. As a first step toward this goal, we identified key cell cycle-regulating genes in the recently published genome of the symbiotic dinoflagellate *Symbiodinium minutum*. We then correlated expression of these genes with cell cycle phase in diurnally growing *S. minutum* in culture. Of particular interest, this approach allowed us to identify cyclins and cyclin-dependent kinases that are involved in the G1/S transition—a likely point for coral cells to exert control over algal cell divisions.

## INTRODUCTION

A critical relationship exists between reef-building corals and the symbiotic algae residing within them. These dinoflagellate algae (zooxanthellae), from the genus *Symbiodinium*, are photosynthetic, harvesting energy from sunlight and sharing that energy with their coral hosts. In return, corals provide them with metabolic intermediates, a stable position in the water column, and protection from grazing. This relationship, as important as it is for reef health, is delicate—exposure to temperatures only marginally above the average summer maximum can cause corals to bleach, expelling their resident algae (reviewed in (Brown 1997)). Massive bleaching due to global warming will drastically and irreversibly alter coral reef ecosystems around the world, adversely impacting fisheries, coastal ecosystems, and placing financial strain on developing economies that depend on tourism (reviewed in (Moberg and Folke 1999)).

Remarkably, studies suggest that failure of the host to properly regulate zooxanthellae cell divisions at high temperatures may cause bleaching (Bhagooli and Hidaka 2002; Strychar *et al*. 2005). In fact, in many corals, the optimal growth temperature for the symbiont exceeds the bleaching threshold. For example, clade C zooxanthellae isolated from *Montipora verrucosa* proliferate better when reared at 31° than when reared at lower temperatures (Kinzie *et al*. 2001); this temperature, however, is lethal to the host (Jokiel and Coles 1977).

To establish symbiosis, corals produce a chemical signal that forces symbionts into a non-motile, dividing state (Koike *et al*. 2004). Unfortunately, it is unclear how corals coordinate host and algal cell divisions to ensure that the proper density of symbionts is maintained. Several pre-mitotic mechanisms have been proposed in a variety of Cnidarians, including factors produced by coral cells that inhibit the algal cell cycle (Smith and Muscatine 1999) and limited access to nutrients (Falkowski *et al*. 1993), which may be controlled by the host or simply by competition among symbionts (consistent with (Mcauley and Darrah 1990)). Postmitotic regulation might include digestion of algae *in situ* (Titlyanov *et al*. 1996) or expulsion of excess algae (Jones and Yellowlees 1997). In the anemone *Aiptasia pulchella*, rates of algal expulsion increase with temperature and expelled algae, surprisingly, have a much higher rate of mitosis than algae that are retained within their hosts (Baghdasarian and Muscatine 2000). Regardless of the mechanism(s) responsible for maintaining a healthy symbiont density, it is evident that controlling algal cell divisions in hospite is important.

The eukaryotic cell cycle is divided into four distinct phases—two gap phases (G1 and G2) interrupted by DNA replication (synthesis/S phase) and followed by mitosis (M), which ends with cytokinesis. Moving successively from one stage to another is governed through phosphorylation of substrates by the cyclin-dependent kinases (CDKs). As this name implies, these proteins are active only when paired with a partner cyclin that can target them to the appropriate substrates. The CDKs are additionally activated by the CDK-activating kinases (CAKs; reviewed in (Kaldis 1999)) and Cdc25 phosphatases and inhibited by members of the WEE1/MYT1 kinase family (reviewed in (Perry and Kornbluth 2007)). Further, the CDK-interacting protein CKS1 is essential for both the G1/S and G2/M transitions (Tang and Reed 1993). Meanwhile, the cyclin-dependent kinase inhibitors (CKIs)—e.g. the mammalian proteins p21^Cip1/Waf1^ and p27^Kip1^—can initiate cell cycle arrest in response to internal or external stimuli (El-Deiry *et al*. 1994; Polyak *et al*. 1994).

While checkpoints regulate each cell cycle transition, eukaryotic cells commit to division during G1. In yeast, this point of commitment is called START (Hartwell *et al*. 1974), while in metazoans it is referred to as the restriction point (R-point) (Pardee 1974). In general, the cell cycle is only sensitive to external factors prior to this; once a cell has committed to divide, it can do so without any further input. In light of this, if a Cnidarian host is regulating the proliferation of its symbionts, it makes sense that it would do so by blocking the R-point. In fact, symbiotic algae in *Aiptasia pulchella* spend considerably longer in G1 than the same species of algae *(Symbiodinium pulchrorum)* in culture, despite the durations of S, G2, and M remaining relatively consistent (Smith and Muscatine 1999).

As a first step toward our goal of understanding how coral cells regulate the divisions of their symbiotic algae, we identified key cell cycle regulators in the recently published genome of *Symbiodinium minutum* (Shoguchi *et al*. 2013). We then correlated the expression of those genes with cell cycle phase in diurnally growing *S. minutum* in culture. This approach allowed us to find specific cyclins and CDKs that are involved in the G1/S transition—a likely point for coral cells to exert control over algal cell divisions.

## MATERIALS AND METHODS

### Identification of Putative Cell Cycle-Regulating Genes

Amino acid sequences for all annotated CDKs in *H. sapiens, A. thaliana*, and *S. cerevisiae* were downloaded from GenBank. An alignment, constructed with Probcons (Do *et al*. 2005), was used to generate a CDK-specific HMMER model (HMMER 3.1b1, hmmer.org). Again using HMMER, this model was queried against the predicted proteome of *S. minutum* ((Shoguchi *et al*. 2013); available at marinegenomics.oist.jp/symb/viewer/info?project_id=21) to identify novel CDKs. Initial results were culled by removing all high scoring pairs (HSPs) with fewer than 200 residues and any HSPs that didn’t begin within 55 bases of the start of the CDK alignment. To identify novel cyclins, Cdc25s, MAT1 homologs, and CKS1 homologs in *S. minutum*, hidden Markov models were retrieved from PFAM (pfam.xfam.org) and queried against the predicted proteome of *S. minutum*, again using HMMER. For the cyclins, a PFAM model describing the characteristic N-terminal domain that is common to all cyclins was chosen. All potential CDK, cyclin, Cdc25, MAT1, and CKS1 sequences were then reciprocally queried against the UniProtKB database using the HMMER web server (Finn *et al*. 2011). Those sequences that had CDKs, cyclins, Cdc25s, MAT1, or CKS1 as top hits were used for further analysis unless they carried additional domains that made them likely members of another protein family (e.g. an N-terminal cyclin domain in a putative CDK).

### Phylogenetic Analysis

Probcons was used to create an alignment of putative *S. minutum* CDKs with representative CDKs from *H. sapiens, A. thaliana*, and *S. cerevisiae*. Gaps were removed from the alignment before generating PhyML (Guindon *et al*. 2010) and BioNJ (Gascuel 1997) trees with 1000 bootstraps that were rooted with a related kinase—MAPK-overlapping kinase isoform 1 (MOK1) from *A. thaliana*. For the cyclins, an alignment of the N-terminal domains of the novel *S. minutum* sequences and representative cyclins from *H. sapiens, A. thaliana*, and *S. cerevisiae* was created using ClustalW ((Larkin *et al*. 2007; Goujon *et al*. 2010); accessed through phylogeny.fr). Gaps were removed from the alignment and trees were constructed as they were for the CDKs.

### Cell Culture

Cultures of S. *minutum* were acquired from the National Center for Marine Algae (Boothbay Harbor, ME; accession number CCMP3345). They were grown in flasks containing silica-free f/2 made from filtered sea water collected from the Gulf of Maine supplemented with 50 mg/ml each of kanamycin, ampicillin, and streptomycin. A period of 13h of light (approximately 100 μmol^*^photons^*^m^2*^s^-1^) followed by 11h of was maintained in an environmental growth chamber set to 25°. Experiments were performed on cultures undergoing log-phase expansion (1-2 x 10^6^ cells/ml).

### Flow Cytometry

To determine the fraction of cultured cells in each cell cycle phase across a 24h day-night cycle, 1 x 10^6^ cells were collected. They were centrifuged for 5 minutes at 200 x g, fixed in 1 ml of 70% ethanol, and resuspended in a staining buffer (PBS, 0.1% Triton X-100, 20 μg/ml propidium iodide (Sigma-Aldrich), and 10 μg/ml RNase A). Cells stained with propidium iodide were run through a FACSCalibur (BD Biosciences, San Jose, CA) set to detect 585 nm fluorescence upon excitation at 488 nm. Data were collected for 10,000 cells and analyzed using FlowJo v.10.1 (Ashland, OR).

### Quantitative RT-PCR

To measure the expression of putative cell cycle regulators in cultured cells across a 24h day-night cycle, 2 x 10^7^ cells were collected by centrifuging at 5,500 x g for 15 min., media was removed, and pellets were flash frozen in liquid nitrogen. RNA extraction combined lysis and homogenization in TRIzol (Life Technologies, Carlsbad, CA) with a kit-based cleanup following the protocol established by Rosic and Hoegh-Guldberg (Rosic and Hoegh-Guldberg 2010). Briefly, pellets were resuspended in 1 mL of TRIzol, mixed thoroughly by pipetting, and transferred to screw-cap tubes with 0.3 g of glass beads (Sigma-Aldrich, St. Louis, MO). Cells were then disrupted twice for 90 sec. in a MagNA Lyser (Roche Life Science, Basel, Switzerland) at 4,500 rpm. Debris was removed by centrifuging at 12,000 x g for 1 min., and then supernatant was run through the RNeasy Plus Mini kit following manufacturer’s instructions (Qiagen, Hilden, Germany). RNA was eluted in 30 μl of nuclease-free water and quantified on a NanoDrop Lite (Thermo Fisher Scientific, Waltham, MA). 1 μg of RNA was reverse transcribed using a QuantiTect Reverse Transcription Kit (Qiagen) following manufacturer’s instructions; this kit includes a genomic DNA wipeout step. cDNA was diluted 1:5 in nuclease-free water before being used for quantitative PCR (qPCR). qPCR reactions were setup as follows: 10 μl QuantiNova SYBR Green PCR Master Mix (Qiagen), 1.5 μl mixed forward and reverse primer (10 μM each, Table S3), 4.5 μl of nuclease-free water, and 4 μl of template. Reactions were run on a CFX96 (Bio-Rad, Hercules, CA) at 95°C for 2 min. for initial activation following by 40 cycles of 95°C for 5 sec. then 60°C for 10 sec. Data were normalized to the expression of S-adenosyl methionine synthase (SAM) (Rosic *et al*. 2011) and analyzed using the 2^-ΔΔCt^ method (Livak and Schmittgen 2001). Average expression levels relative to the 08:00 time-point were clustered using MeV 4.9.0.

### Data Availability

File S1 contains flow cytometry data (FCS files) for Figure 1. Individual FCS files are ordered by time-point (three files for each time-point from 0:00 to 20:00). File S2 contains all real-time RT-PCR analysis with a separate tab for each gene.

**Figure 1:**
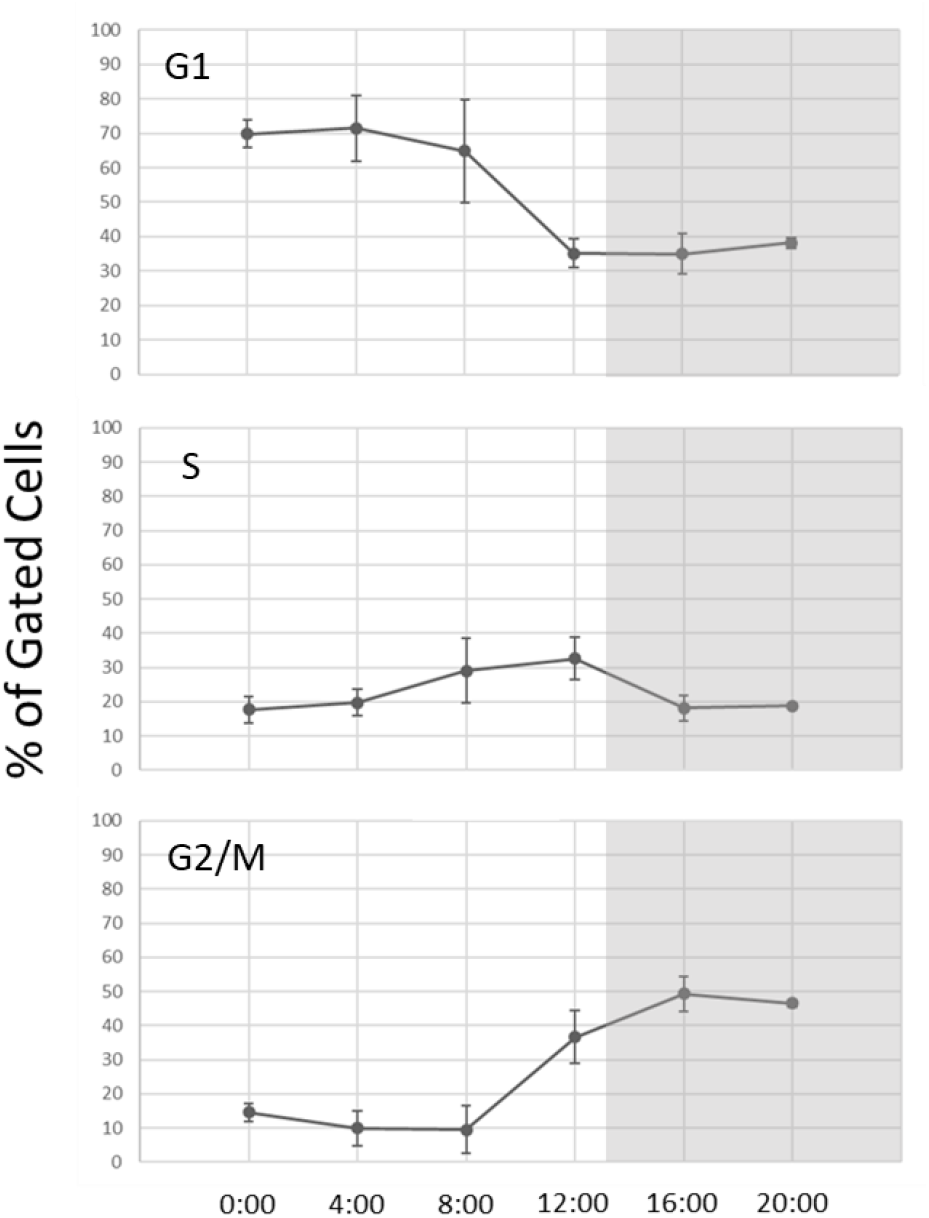
Cell cycle distribution of free-living *S. minutum* over 24h. *S. minutum* maintained on a diurnal cycle were collected from a culture every 4h. Cells were stained with propidium iodide and nuclear content was assessed by flow cytometry. Data were analyzed using the FlowJo Cell Cycle platform and plotted as the average percentage of gated cells in each cell cycle phase +/-standard deviation (n = 3). Data are representative of multiple biological replicates. Grey background represents periods of darkness.

## RESULTS AND DISCUSSION

Amino acid sequences for annotated cell cycle regulators (belonging to the following families: CDKs, cyclins, CAKs, CKS1, WEE1/MYT1, Cdc25, and CKIs) in *H. sapiens, A. thaliana*, and *S. cerevisiae* were used to identify homologous sequences in the genome of *S. minutum*. The expression of these novel cell cycle genes was monitored throughout the cell cycle to help validate their function as predicted by homology.

Division of free-living *Symbiodinium* is regulated in response to light, so cells grown on a diurnal cycle (13h of light to 11h of darkness) were sampled every 4h over one 24h period to determine the percentage of cells in each cell cycle phase at a given time. Cells were stained with the nuclear stain propidium iodide and nuclear content was assessed by flow cytometry. The highest percentage of G1 cells—about 70%—was observed at the onset of the light period (time 00:00) and persisted until the onset of DNA replication at 04:00 (Figure 1, Figure S1). By 08:00, the fraction of G2 cells began to rise later as the fraction of G1 cells fell. S phase peaked around the initiation of the dark period (13:00), with the largest percentage of G2 cells and lowest percentage of G1 cells observed a few hours later. Cytokinesis occurred in the hours immediately prior to the start of the light period. These results are consistent with published data, which show that the vast majority of free-living *Symbiodinium* are in G1 early in the day and that the proportion of G2 cells peaks during the dark period (Smith and Muscatine 1999; Wang *et al*. 2013). It is worth noting that our results indicate a background presence of S phase cells (about 10%) at all time points. Most likely, this background is an artifact of the modeling that was done to fit a histogram of DNA content to specific cell cycle phases in FlowJo. These cells probably belong to either the G1 or G2 population.

When attempting to assign molecular events, such as changes in gene expression, to a particular cell cycle phase, it is commonplace to use a synchronized cell population; this is often achieved by treating with a mitosis inhibitor such as nocodazole. While this is generally desirable, we chose not to use a synchronized population. First of all, nocodazole is cytotoxic and is known to affect gene expression (Cho *et al*. 2006). Second, while darkness leads to the accumulation of cultured *Symbiodinium* cells in G1, we were unable to increase the percentage of G1 cells with prolonged darkness (unpublished data). Still, with a large fraction of cells entering S phase at 04:00, we would expect to see increased expression of genes involved in the G1/S transition at that time. Likewise, the proportion of G2 cells drops precipitously within the last 4 hours of darkness, so we would expect to see increased expression of genes involved in mitosis during this period.

### CDKs

Nine CDKs were identified in *S. minutum;* to group them within accepted subclasses (e.g. CDKA, CDKB), phylogenetic trees were constructed using both maximum likelihood (PhyML) and neighbor joining (BioNJ) methods. Both methods yielded similar topologies, so the BioNJ tree was chosen for publication. Of the nine CDKs, two clustered with the CDKA/CDK1-3 family (Figure 2). CDKs from this family all contain the highly conserved PSTAIRE motif that seems to be important for their association with the cyclins that are most responsible for cell cycle progression (Figure 3A). Further supporting their role in cell cycle regulation, the mRNA expression for both of these *S. minutum* CDKs is highest from 04:00 to 16:00 (Figure 4). It is precisely during this time that we observe an increase in the percentage of S phase cells (starting at 04:00) and subsequently G2 cells (starting at 08:00) (Figure 1). Based on these data, and sticking with the nomenclature established for cell cycle genes in other protists (e.g. (Huysman *et al*. 2010)), we are confident in classifying these genes as *S. minutum* CDKA1 and CDKA2. Interestingly, mRNA expression for CDKA1 begins to increase earlier than the mRNA expression for CDKA2, suggesting that CDKA1 may be important for S phase entry and CDKA2 for progression into G2.

**Figure 2:**
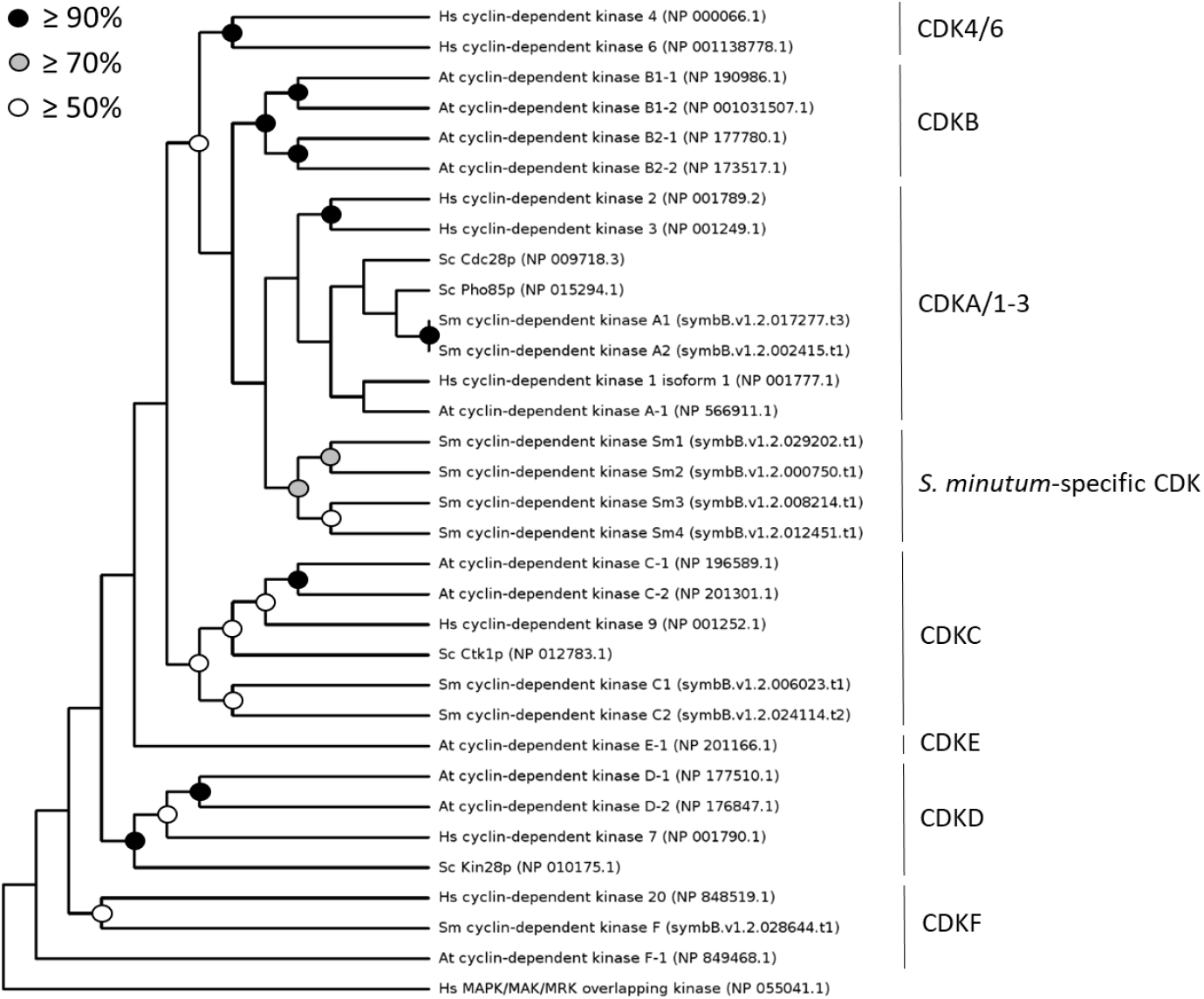
Phylogenetic analysis of the CDKs of *S. minutum*. Protein sequences for 9 CDKs identified in the genome of *S. minutum* (labelled Sm) were aligned with representative CDK sequences from *H. sapiens* (Hs), *A. thaliana* (At), and *S. cerevisiae* (Sc) using Probcons. Phylogenetic trees created using either neighbor-joining (BioNJ) or maximum likelihood (PhyML) methods yielded similar topologies. The tree shown here was constructed using BioNJ with 1,000 replicates. NCBI accession numbers or specific genome identifiers (for *S. minutum)* are indicated next to the sequence ID.

**Figure 3:**
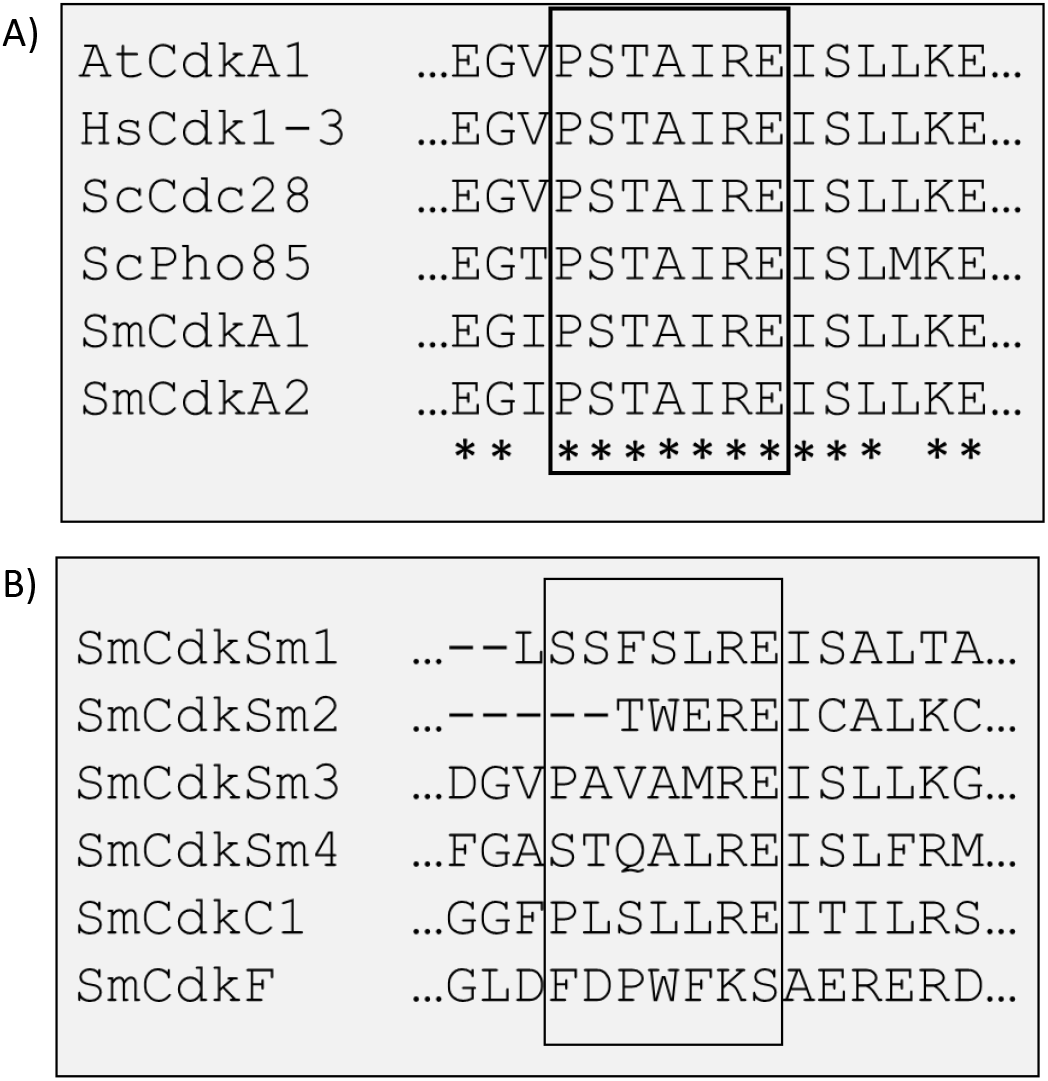
The cyclin binding motif in the CDK family. Protein sequences for the CDKA/1-3 family members from *H. sapiens* (Hs), *A. thaliana* (At), and *S. cerevisiae* (Sc) were aligned to the CDK protein sequences identified in *S. minutum* (Sm) using ClustalW. Sixteen residues flanking the cyclin binding motif are shown. Conserved amino acids are marked by an asterix in the bottom row. **A)** CDKA/1-3 family members from each species contain a highly conserved cyclin binding motif (PSTAIRE). **B)** Comparison of the cyclin binding motif in all *Symbiodinium* CDKs.

**Figure 4:**
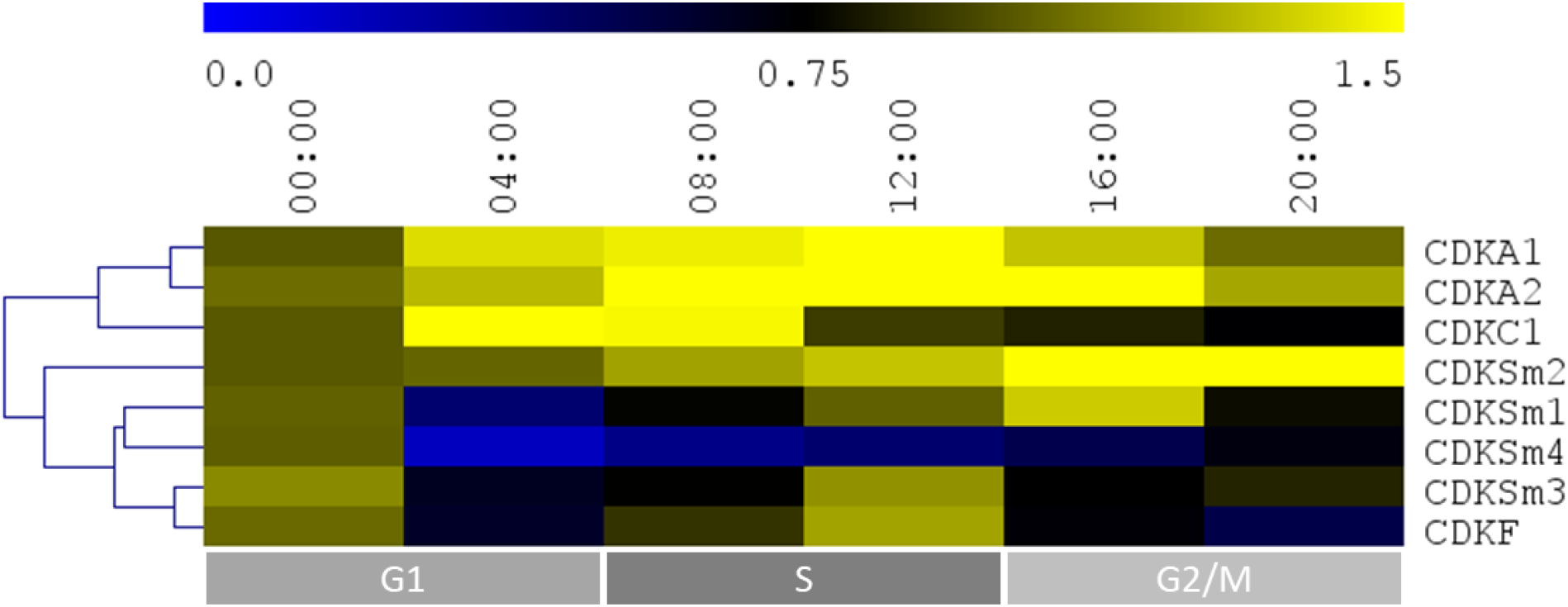
Hierarchical clustering of the gene expression profiles of CDKs in *S. minutum*. Messenger RNAs were isolated from free-living *S. minutum* maintained on a diurnal cycle every 4h. Following reverse transcription, qPCR was run with primers for specific CDKs. Data were normalized to the expression of SAM and analyzed using the 2^-ΔΔCt^ method. Average expression levels relative to the 0:00 time-point were clustered using MeV 4.9.0 (n = 3). Data are representative of multiple biological replicates. Grey boxes below indicate the predominant cell cycle phase at a given time-point.

In addition to the two CDKAs, our phylogenetic analysis revealed two CDKCs, a CDKF that clustered with human CDK20, and a cluster of four *S. minutum*-specific CDKs (Figure 2). The CDKC family, including CDK9 in animals and Ctk1p in *S. cerevisiae*, is involved in transcription initiation (Wallenfang and Seydoux 2002). Of the two CDKCs identified computationally, only one of them, which we call CDKC1, had detectable mRNA expression at any point over a 24h time-course (Figure 4), with expression increasing immediately after the onset of the light period and remaining high through at least 08:00. This expression probably reflects a general increase in gene expression as cells prepare to enter S phase and mitosis. As in other eukaryotes, expression of CDKC1 is not likely to depend on cell cycle phase. As for the CDKF homolog, its expression peaks at or near the end of the light period before dropping continuously through the dark hours (Figure 4). Interestingly, in the green algae, *Chlamydomonas reinhardtii*, a CDKF homolog is critical for assembly of flagella (Tam *et al*. 2007). In *Symbiodinium*, it has been established that a non-motile dividing cell, lacking flagella, will give rise to two motile daughter cells (Fitt and Trench 1983). The highest fraction of motile cells is observed within the first few hours of the light period (Fitt and Trench 1983), which can be explained by the fact that cytokinesis occurs within the last few hours of darkness (Figure 1). Increased expression of CDKF as cells enter G2 may indicate that the daughter flagella in *S. minutum* are assembled prior to mitosis; this would be an interesting area for further investigation. Finally, the *S. minutum*-specific CDKs (CDKSm1-4) are each expressed differently throughout the day (Figure 4). CDKSm1 and CDKSm2, for instance, reach peak expression during the dark period, suggesting they may have a role in preparing the cell for mitosis. Interestingly, CDKSm4 has minimal expression except for the 4h immediately prior to the onset of the light period—the time during which mitosis occurs. Finally, CDKSm3 has a similar expression pattern to CDKF; its expression peaks at or near the end of the light period then decreases through the dark hours. It is possible that CDKSm3 partners with CDKF in flagellar assembly or plays some other role critical during G2.

### Cyclins

Fifteen cyclins were identified in *S. minutum*. From the N-terminal domain that is present in all cyclins, phylogenetic trees were constructed using both maximum likelihood (PhyML) and neighbor joining (BioNJ) methods. Both trees had similar topologies, so the BioNJ tree was chosen for publication. Based on this phylogeny, the *S. minutum* cyclins were grouped within accepted subclasses (e.g. cyclin A, cyclin B). This grouping did not reveal any sequences with a high degree of similarity to known cyclin A family members (Figure 5). Analysis of gene expression, however, revealed that a particular cyclin transcript (identified as cyclin A) reaches its peak expression at 12:00 (Figure 6). During this time the percentage of S phase cells also peaks (Figure 1). It is known that A-type cyclins are active in S phase and are, in fact, necessary to initiate DNA replication (Girard *et al*. 1991). Further supporting our identification of a cyclin A homolog in *S. minutum*, hierarchical clustering of gene expression indicated that its expression most closely matches the expression of CDKA1—the CDK that appears to be involved in S phase entry (Figure S2). Additionally, three cyclin B family members were identified (Figure 5). B-type cyclins are the “mitotic cyclin” that is necessary for proper spindle assembly (Michel *et al*. 2004) and whose rapid degradation precipitates anaphase (Holloway *et al*. 1993). As such, we would expect to see high levels of cyclin B mRNA from 16:00 to 20:00 (i.e. when the highest percentage of cells is in G2 or mitosis) (Figure 1). This, however, is not the case; mRNA expression of both cyclin B1 and B3 peaks around 12:00, while cyclin B2 transcript remains steady throughout the day before dropping off during the dark hours (Figure 6). It is possible that the bulk of transcription in *Symbiodinium* occurs during the daylight hours, though this is certainly not the case for all transcripts we investigated. Another possibility is that while expression of the B-type cyclins drops off before the majority of cells make it into G2, the protein has accumulated to sufficiently high levels and will remain stable until the cells are ready to complete mitosis. Undoubtedly, post-translational control of cyclins is critical for proper coordination of the cell cycle. For example, the ubiquitin proteasome pathway is critical for causing the rapid degradation of cyclin B that triggers anaphase (Holloway *et al*. 1993). Oddly, no cyclin D homolog was uncovered. This family of cyclins is involved in bringing cells out of quiescence and driving them through G1 (Resnitzky *et al*. 1994). Cyclin D appears to have arisen early in the eukaryotes, so the lack of a homolog in *S. minutum* likely reflects a deletion somewhere in its lineage (Cao *et al*. 2014). Similarly, no cyclin E homolog was detected in *S. minutum*. Initial work suggested that cyclin E is absent in protists (Ma *et al*. 2013), however, analysis of several unicellular choanoflagellates uncovered cyclin E homologs (Cao *et al*. 2014). Cyclin E seems to have arisen in the lineage leading to choanoflagellates and animals after splitting from the lineage that led to dinoflagellates.

**Figure 5:**
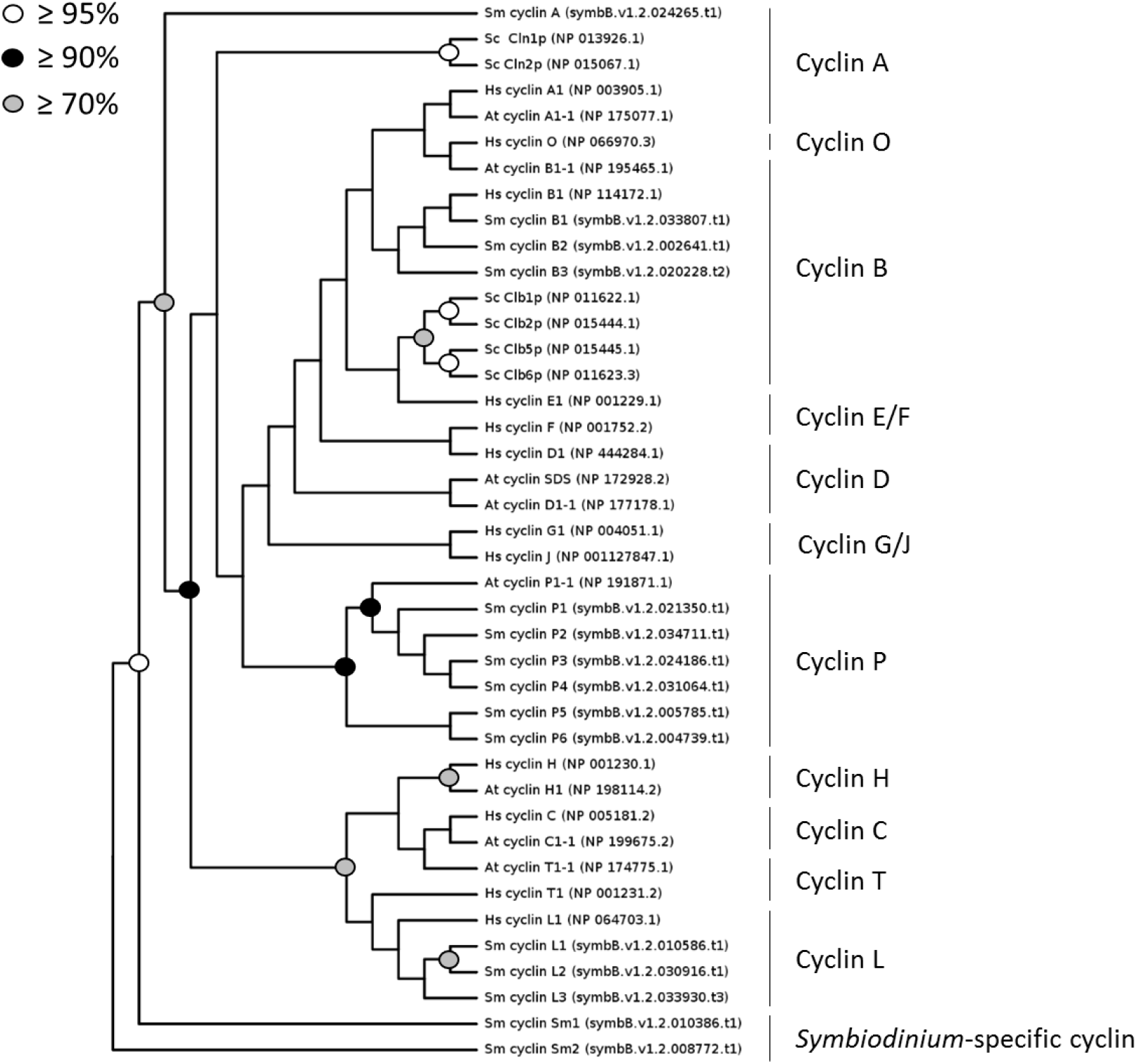
Phylogenetic analysis of the cyclins of *S. minutum*. Protein sequences of the N-terminal domains for 15 cyclins identified in the genome of *S. minutum* (labelled Sm) were aligned with representative cyclin N-terminal domain sequences from *H. sapiens* (Hs), *A. thaliana* (At), and *S. cerevisiae* (Sc) using ClustalW. Phylogenetic trees created using either neighbor-joining (BioNJ) or maximum likelihood (PhyML) methods yielded similar topologies. The tree shown here was constructed using BioNJ with 1,000 replicates. NCBI accession numbers or specific genome identifiers (for *S. minutum)* are indicated next to the sequence ID.

**Figure 6:**
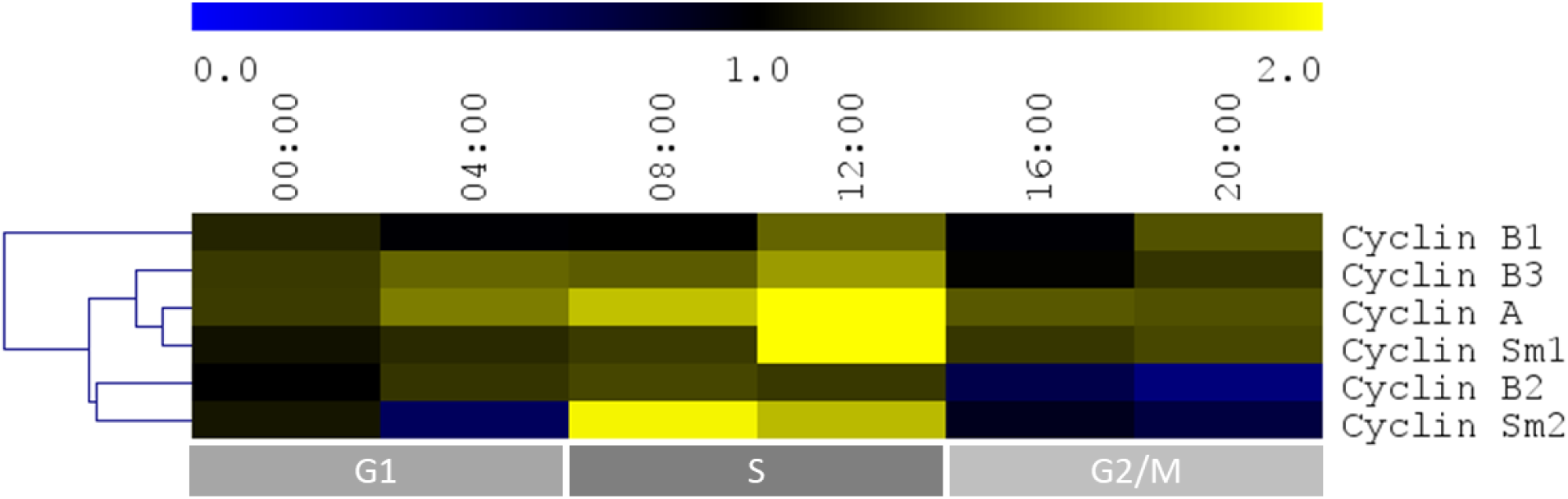
Hierarchical clustering of the gene expression profiles of cyclins in *S. minutum*. Messenger RNAs were isolated from free-living *S. minutum* maintained on a diurnal cycle every 4 hr. Following reverse transcription, qPCR was run with primers for specific cyclins. Data were normalized to the expression of SAM and analyzed using the 2^-ΔΔCt^ method. Average expression levels relative to the 0:00 time-point were clustered using MeV 4.9.0 (n = 3). Data are representative of multiple biological replicates. Grey boxes below indicate the predominant cell cycle phase at a given time-point.

In addition to these canonical cell cycle-regulating cyclins, we identified six P-type cyclins, three L-type cyclins, and two putative cyclins that do not fit phylogenetically into the well-established families (Figure 5). A reciprocal query of the two unidentified sequences against the UniProtKB database using the HMMER web server yielded no clues about their identity; nevertheless, a variety of cyclin sequences were returned as top hits, leading us to classify these as *Symbiodinium*-specific cyclins. Both—cyclin Sm1 and Sm2—are expressed in a seemingly cell cycle-specific way. Cyclin Sm1 is expressed similarly to cyclin A, peaking at 12:00 (Figure 6). Similarly, cyclin Sm2 is at its highest level of expression from 08:00 to 12:00 (Figure 6). During this time, cells go through S phase (Figure 1), suggesting roles for cyclins Sm1 and Sm2 in DNA replication. The six P-type cyclins are particularly interesting because cyclins belonging to this family are thought to be involved in phosphate signaling (Torres Acosta *et al*. 2004). Such a link connecting phosphate signaling to cell cycle regulation would be unsurprising in *Symbiodinium* since concurrent nitrogen and phosphate limitation can induce G1 arrest in these algae (Smith and Muscatine 1999). Under the conditions we tested, the cyclins P1, P2, and P3 have relatively steady expression across 24h (Figure 7). Cyclin P4 is present at a level that is too low for analysis, while cyclin P5 is undetectable. Since inorganic phosphate is abundant in f/2 media, steady expression of these P-type cyclins is unsurprising. Cyclin P6, however, has an interesting expression pattern, peaking from 04:00 to 08:00 before decreasing substantially. It is during this time that cells are transitioning from G1 to S phase. Perhaps cyclin P6 is functioning as a nutrient sensor that allows cells to enter S phase if there is abundant phosphate. Finally, the L-type cyclins are important for RNA splicing (Dickinson *et al*. 2002). Transcripts for the three L-type cyclins that were identified computationally, however, were undetectable.

**Figure 7:**
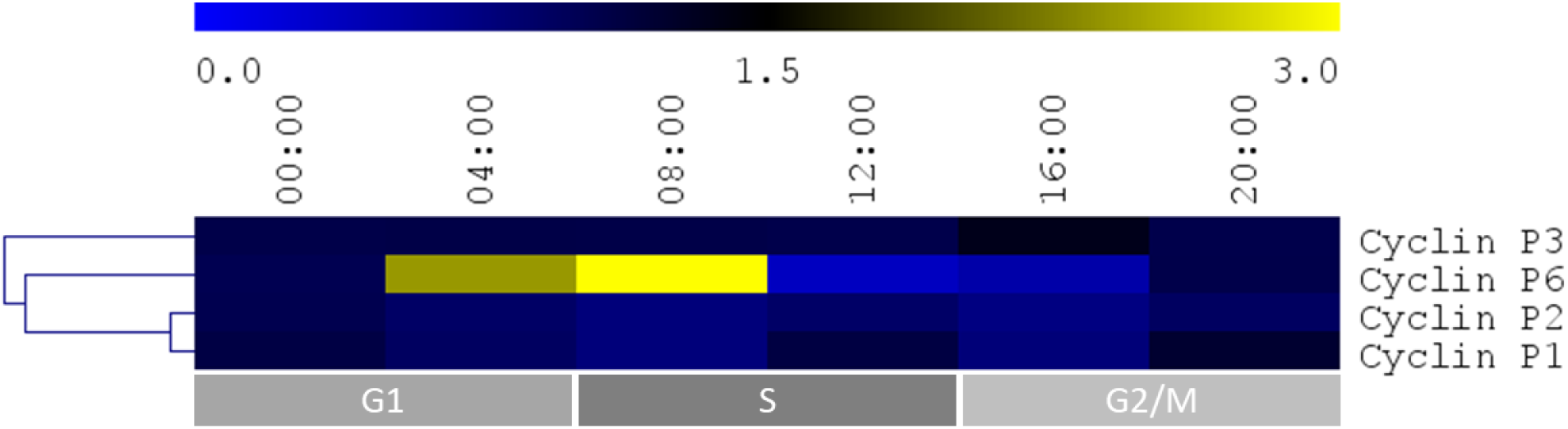
Hierarchical clustering of the gene expression profiles of P-type cyclins in *S. minutum*. Messenger RNAs were isolated from free-living *S. minutum* maintained on a diurnal cycle every 4 hr. Following reverse transcription, qPCR was run for specific cyclins P family members. Data were normalized to the expression of SAM and analyzed using the 2^-ΔΔCt^ method. Average expression levels relative to the 0:00 time-point were clustered using MeV 4.9.0 (n = 3). Data are representative of multiple biological replicates. Grey boxes below indicate the predominant cell cycle phase at a given time-point.

### CDK Activators and Inhibitors

In addition to the presence or absence of a cyclin partner, CDKs are regulated by numerous other modulators. The CAKs and Cdc25 phosphatases, for instance, work in concert to add activating phosphorylations and remove inhibitory phosphorylations from CDKs (reviewed in (Kaldis 1999; Perry and Kornbluth 2007). In contrast, members of the WEE1/MYT1 kinase family phosphorylate CDKs in a way that renders them inactive (reviewed in (Perry and Kornbluth 2007). Further regulating CDK activity, the CDK-interacting protein CKS1 functions as a docking protein that allows CDKs to interact with their substrates (Tang and Reed 1993). Finally, the CKIs can block CDK activity and initiate cell cycle arrest in response to internal or external stimuli (El-Deiry *et al*. 1994; Polyak *et al*. 1994).

From this list of CDK activators and inhibitors, only a partial CAK subunit and three CKS1 homologs were identified in S. *minutum*. The CAK subunit was identified as MAT1—one part of a trimeric CAK complex (reviewed in (Kaldis 1999)). Surprisingly, however, only the N-terminus of this MAT1 homolog aligned with MAT1 sequences from other species. For example, MAT1 in *H. sapiens* is 309 amino acids long but only the first 140 residues aligned with the *S. minutum* protein, which itself is 245 amino acids long. The transcript for this partial MAT1 is expressed at relatively steady levels throughout the day, but drops just before the onset of the dark period (Figure 8). Canonically, MAT1 functions as an assembly factor, forming a CAK with cyclin H and a D-type CDK. Since there are no cyclin H or CDKD homologs in *S. minutum*, it is likely that this partial MAT1 plays another role, perhaps independent of the cell cycle. The evolutionary history of this sequence might hold clues to its function. As for the three CKS1 homologs, only two— CKS1A and CKS1B—had detectable transcripts (Figure 8). The expression of CKS1B is highest just before darkness and has a similar expression pattern to cyclin B1 (Figure S2), perhaps indicating that these two bind together to a target CDK. Expression of CKS1A is less predictable across the cell cycle.

**Figure 8:**
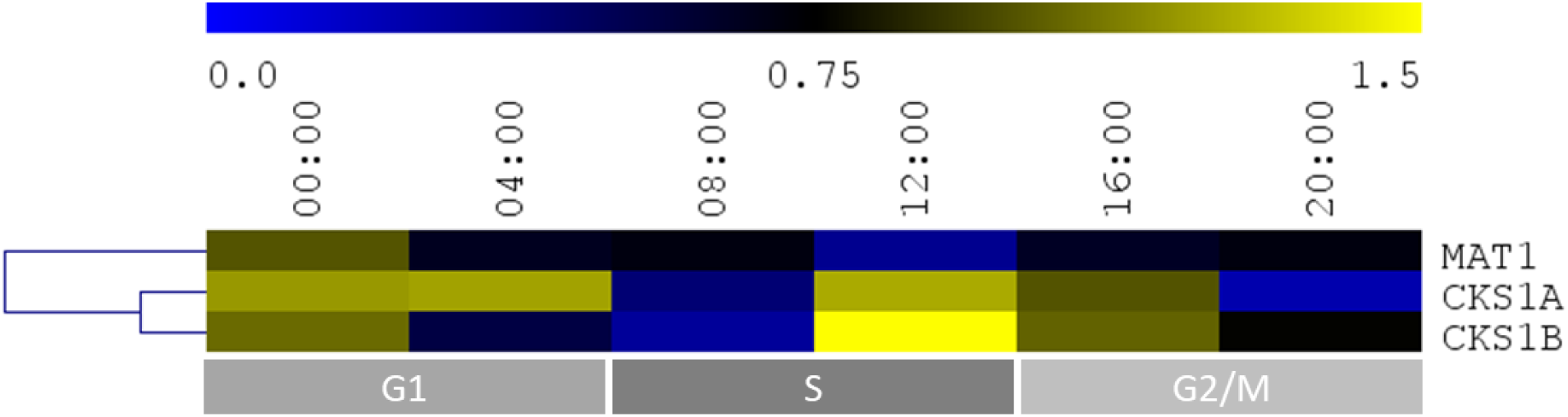
Hierarchical clustering of the gene expression profiles of cyclin/CDK interacting proteins in *S. minutum*. Messenger RNAs were isolated from free-living *S. minutum* maintained on a diurnal cycle every 4 hr. Following reverse transcription, qPCR was run with primers for specific cyclins. Data were normalized to the expression of SAM and analyzed using the 2^-ΔΔCt^ method. Average expression levels relative to the 0:00 time-point were clustered using MeV 4.9.0 (n = 3). Data are representative of multiple biological replicates. Grey boxes below indicate the predominant cell cycle phase at a given time-point.

No members of the WEE1/MYT1 family of CDK-inhibiting kinases were identified; thus, it is not surprising that their antagonists—the Cdc25 phosphatases—were also absent. Finally, no CKIs—small CDK inhibitors such as the mammalian proteins p21^Cip1/Waf1^ and p27^Kip1^—were uncovered in the *S. minutum* genome. It is possible that one or more CKIs are present but are simply undetectable using our methods, which rely on a large degree of sequence similarity that is uncharacteristic of the CKIs.

### Conclusion

Unsurprisingly, our analysis of key cell cycle regulators in the genome of *S. minutum* has revealed that control of mitotic cell divisions in these dinoflagellates shares many features with other eukaryotes. For instance, the two CDKAs that were identified both contain a highly conserved PSTAIRE motif, suggesting that they regulate cell cycle progression directly. Further, CDKA1 has a similar expression profile to cyclin A, suggesting that these proteins form a cyclin/CDK pair that may be involved in driving cells through S phase, much like cyclin A/CDK2 in mammals. Similarly, three cyclin B family members were identified that may pair with one or both CDKAs and play a role in G2 and mitosis—exactly what happens in the mammalian cell cycle. Knowing the identity of these cell cycle regulators will allow us to design experiments to determine the molecular events that are responsible for maintaining Cnidarian control over symbiont life cycles. The cyclin A/CDKA1 combination is particularly interesting because it seems to be the pair that is necessary for initiating S phase. Eukaryotic cells commit to division at a point in G1 or at the G1/S transition. As a general rule, the cell cycle is only sensitive to external factors prior to this commitment point (Pardee 1974; Hartwell *et al*. 1974). Thus, if a Cnidarian host is regulating cell divisions in its endosymbiont, it is likely that it does so by controlling the G1/S transition that seems to be regulated by the cyclin A/CDKA1 pair.

## ACKNOWLEDGEMENTS

Special thanks to Natalie Spivey for critical review of this manuscript.

---

**Figure S1:**
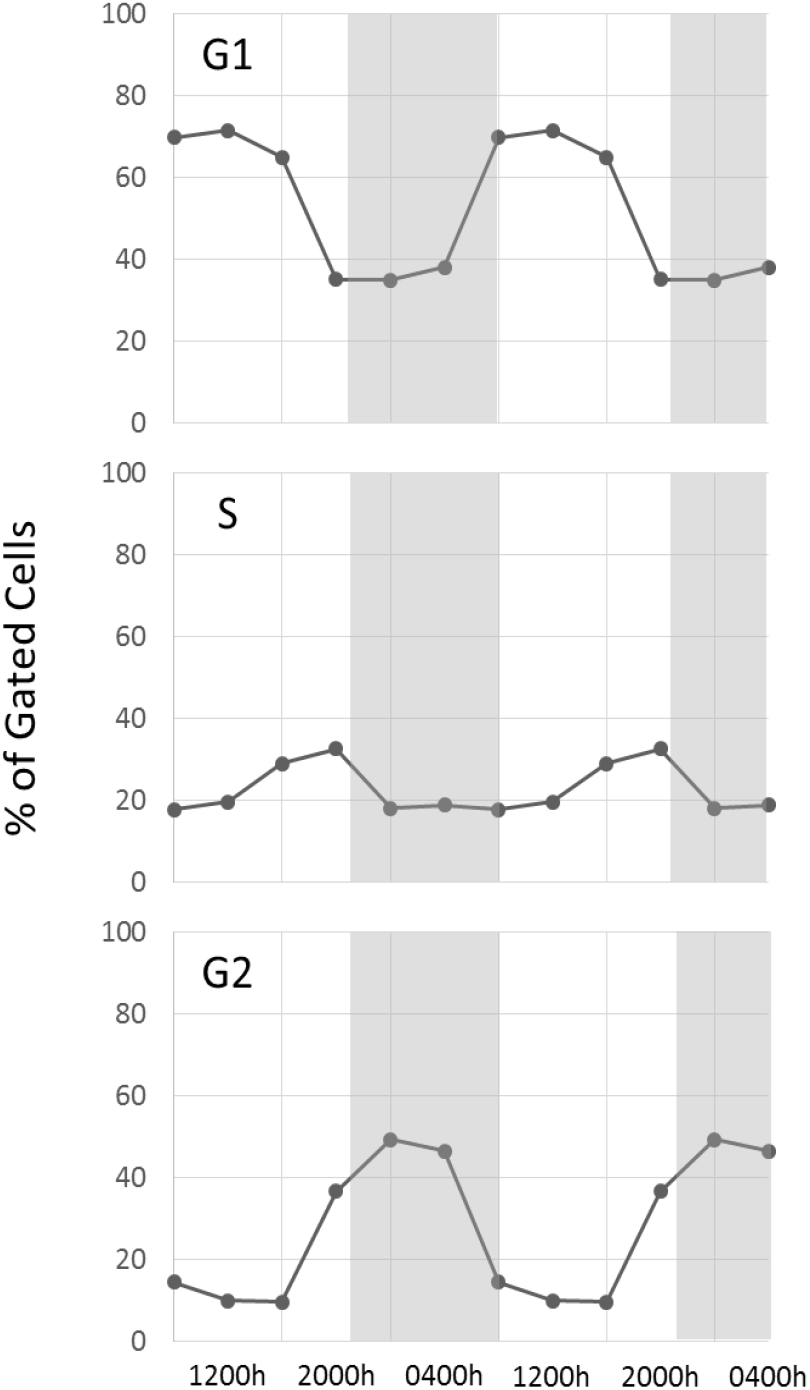
Cell cycles distribution of free-living *S. minutum*. These data were collected over 24h as described for Figure 1. Here, however, the data are duplicated for a second 24h period to show cyclical cell divisions over 48h. Grey background represents periods of darkness.

**Figure S2:**
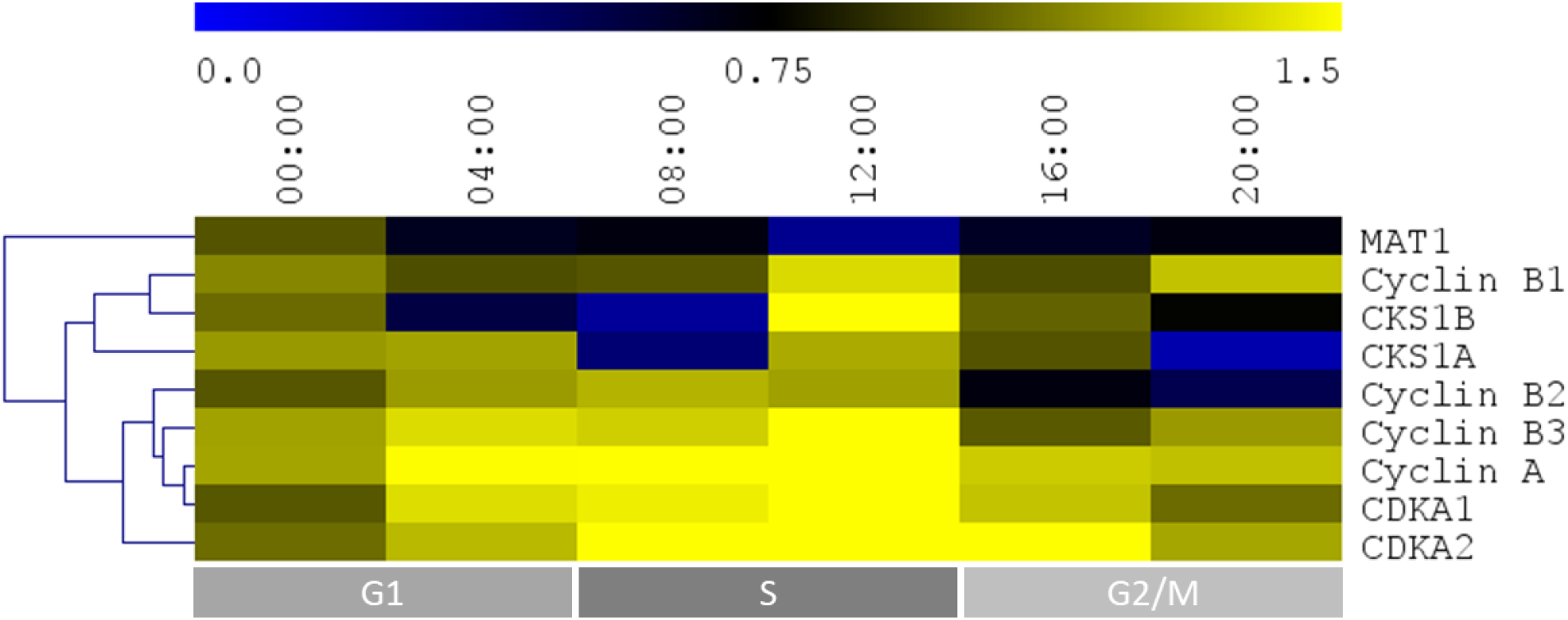
Hierarchical clustering of the gene expression profiles of putative cell cycle regulators in *S. minutum*. Messenger RNAs were isolated from free-living *S. minutum* maintained on a diurnal cycle every 4h. Following reverse transcription, qPCR was run with primers for specific cyclins. Data were normalized to the expression of SAM and analyzed using the 2^-ΔΔCt^ method. Average expression levels relative to the 0:00 time-point were clustered using MeV 4.9.0 (n = 3). Data are representative of multiple biological replicates. Grey boxes below indicate the predominant cell cycle phase at a given time-point.

**Figure S3:**
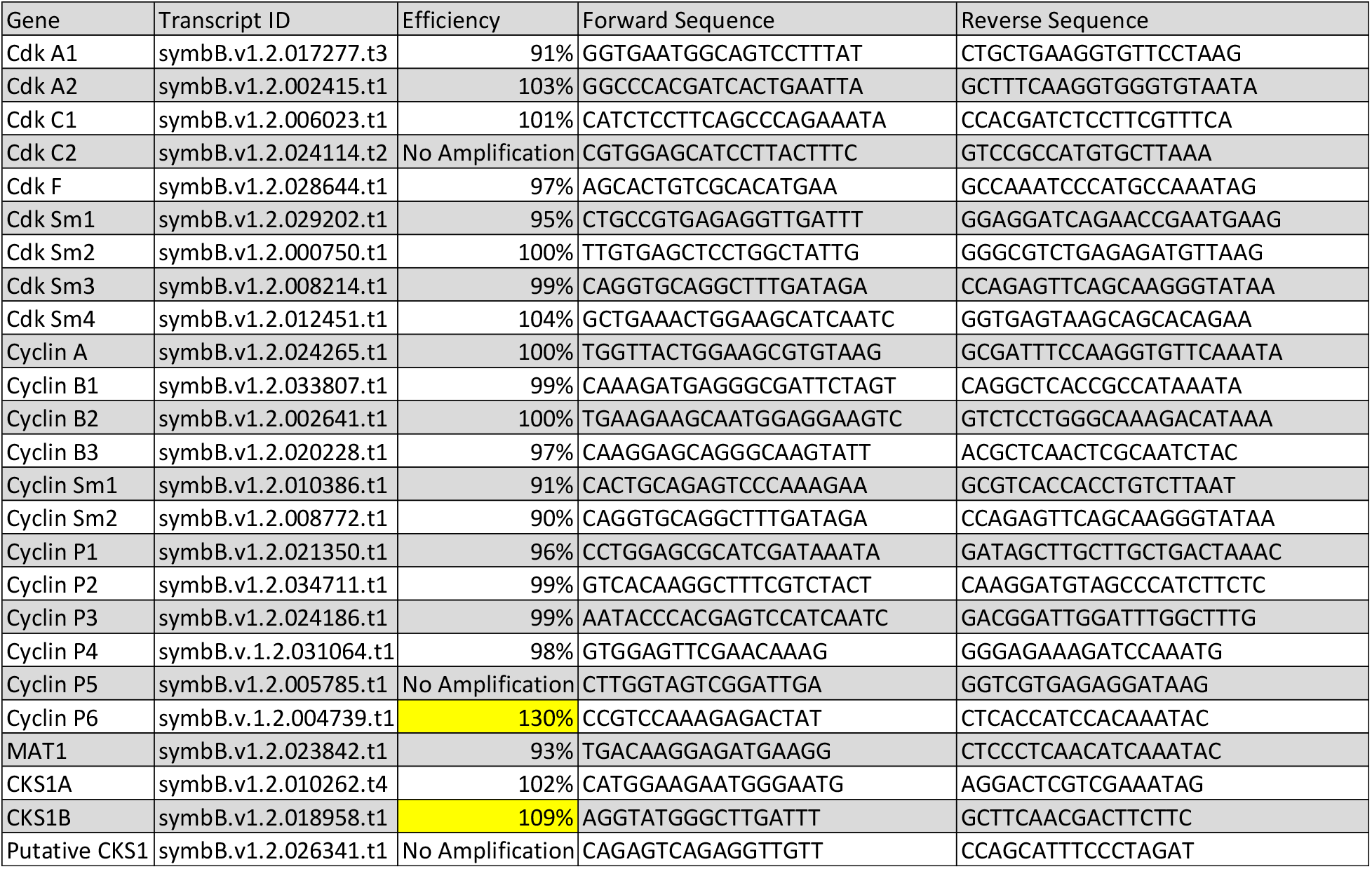
Real-time RT-PCR primers used to amplify potential cell cycle genes in *S. minutum*. Primer sets with efficiency values between 90% and 105% were considered to be useful assays.

